# Collateral consequences of oxidative stress responses result in mitomycin C sensitivity

**DOI:** 10.64898/2026.07.29.741606

**Authors:** Kubra Yigit, Anna Chapman, Peter Chien

## Abstract

Bacterial survival depends on carefully balanced antioxidant defenses against reactive oxygen species. *Caulobacter crescentus* employs the transcription factor OxyR to activate hydrogen peroxide detoxification genes, a seemingly straightforward protective strategy. Here, we reveal an unexpected consequence of environment or genetic activation of peroxide resistance via OxyR resulting in vulnerability to reductively activated antibiotics. An activated OxyR allele protects against peroxide stress due to upregulation of the catalase-peroxidase KatG but simultaneous induction of the AhpCF reductase sensitizes cells to the reductively activated genotoxin mitomycin C. We show that this collateral vulnerability can be induced by brief exposure to oxidative stress and extends to bacteria beyond *Caulobacter*. Our findings illuminate a hidden cost of antioxidant signaling as robust defense against oxidative stress paradoxically creates exploitable vulnerabilities to chemotherapeutic prodrugs.

**SIGNIFICANCE:** Cellular stress responses are vital for survival, but our work reveals an underappreciated principle where activation creates trade-offs with far-reaching consequences. Using *Caulobacter crescentus*, we show that strengthened antioxidant defenses against peroxide simultaneously increase vulnerability to mitomycin C. Because these regulatory mechanisms are widespread across bacteria, our findings suggest a general model wherein stress-defense pathways unavoidably compromise resistance to alternative threats. This conceptual framework not only advances our understanding of stress-response regulation but also identifies new strategies for therapeutic exploitation of bacterial vulnerabilities.

## INTRODUCTION

Bacteria encounter oxidative stress throughout their life cycle from both intracellular metabolic processes and environmental sources^1^. These stressors generate reactive oxygen species (ROS) including superoxide anions, hydrogen peroxide, and hydroxyl radicals that damage cellular DNA^2,3^, proteins^4–6^, and lipids^7^. To survive, cells have evolved sophisticated antioxidant response pathways that detect, neutralize, and scavenge ROS while suppressing their production and mitigating damage^8^. A defining feature of bacterial antioxidant defense is transcriptional regulation by redox-sensitive transcription factors such as PerR, SoxRS, and OxyR. These factors sense changes in cellular redox status and reprogram gene expression accordingly^9^.

OxyR stands out as a central positive regulator of antioxidant genes across many bacterial species and is important for hydrogen peroxide (H_2_O_2_) resistance^10–14^. OxyR operates as a molecular redox sensor where conserved cysteine residues remain as thiols under reducing conditions but form an intramolecular disulfide bond when oxidized by H_2_O ^15,16^. This redox-dependent switch triggers a conformational change that enables OxyR to bind target promoters and activate key antioxidant enzymes, including catalases and peroxidases^12^.

For *Caulobacter crescentus*, an oligotrophic bacterium inhabiting freshwater and soil^15–17^, oxidative stress is a persistent challenge. As an obligate aerobe, *Caulobacter* continuously generates ROS as respiratory byproducts while facing external stressors such as UV radiation and redox-active compounds^1^. *Caulobacter* manages this burden through an antioxidant defense network largely controlled by OxyR^14^. Critical targets include *katG*, encoding catalase-peroxidase KatG, and *ahpC* and *ahpF*, encoding the subunits of alkyl hydroperoxide reductase AhpCF. Together, these enzymes maintain redox homeostasis and protect against oxidative damage.

During our investigation of the oxidative stress response in *Caulobacter crescentus*, we identified a point mutation that hyperactivates OxyR and exposes a physiological trade-off. Although the mutant exhibits enhanced tolerance to H_2_O_2_ and tert-butyl hydroperoxide (TBHP), it suffers reduced fitness under non-stress conditions and heightened susceptibility to the DNA-damaging agent mitomycin C (MMC). RNA-seq reveals an elevated expression of canonical OxyR targets and subsequent deletion analysis shows that KatG primarily drives H_2_O_2_ resistance, while AhpCF underlies sensitivity to MMC, which requires enzymatic reduction to be fully active and similar effects were seen with other prodrugs. Notably, brief H_2_O_2_-induced antioxidant responses were sufficient to increase AhpCF-dependent MMC sensitivity in wild type *Caulobacter* and similar effects were observed in *Escherichia coli*. These findings suggest an interplay between oxidative and reductive stress responses and raise the possibility that precise regulation of antioxidant pathways is essential for survival when cells are exposed to multiple stresses.

## RESULTS

### A point mutation in *oxyR* confers enhanced H_2_O_2_ resistance

To identify genetic determinants of H_2_O_2_ tolerance in *Caulobacter crescentus*, we screened for spontaneous suppressors with increased resistance. Whole-genome sequencing of an isolate with markedly elevated H_2_O_2_ tolerance revealed a point mutation in *oxyR* (CCNA_03811) predicted to cause an A105V substitution. Reconstruction of this allele (oxyR*) in wild type background recapitulated the phenotype where the OxyR* strain exhibited increased resistance to both H_2_O_2_ and TBHP, while sensitivity to the superoxide generator paraquat remained unchanged (Figure 1A, B; Figure S1).

**Figure 1.**
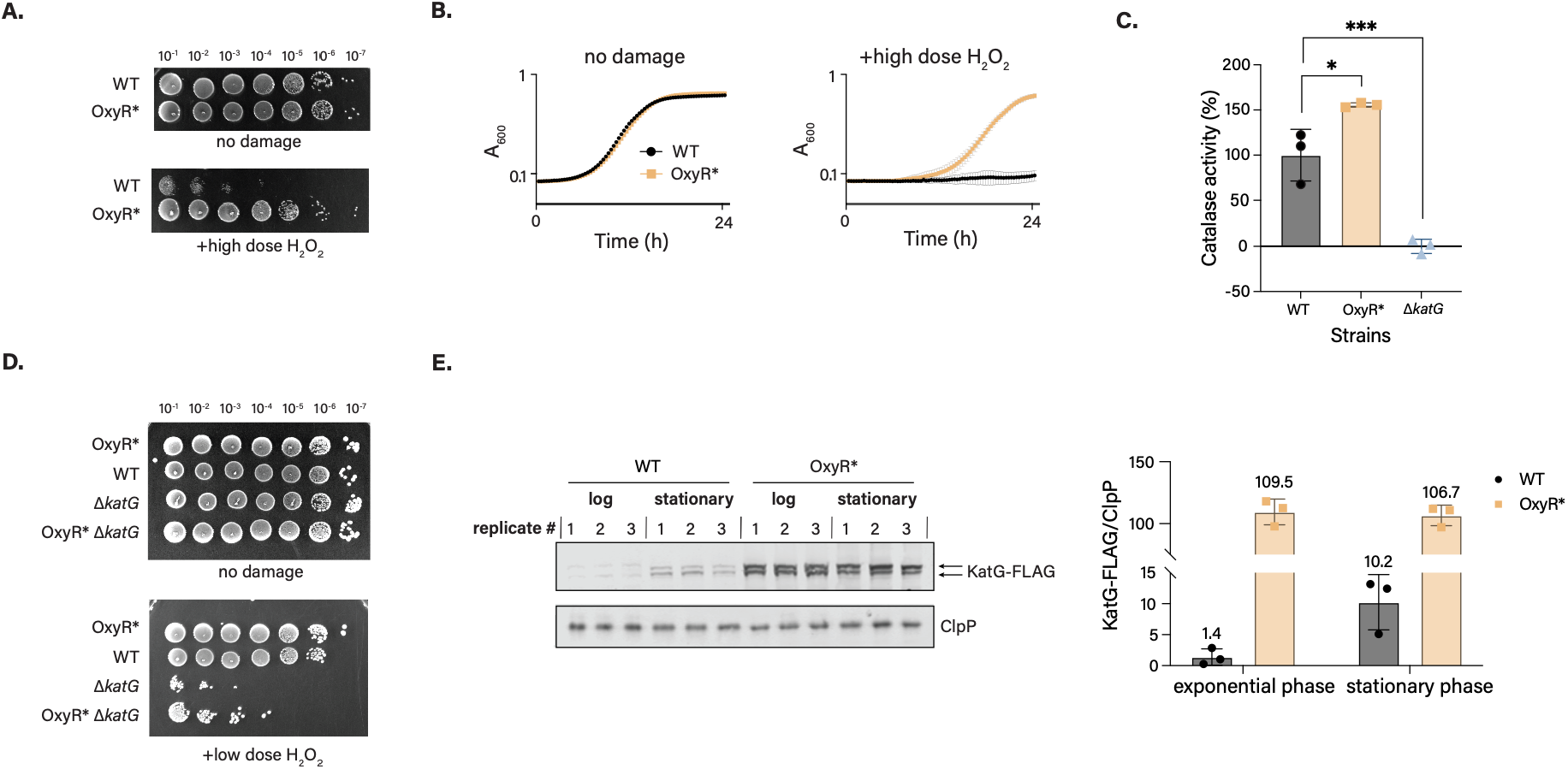
OxyR* strain has increased resistance to H_2_O_2_ due to constitutive *katG* induction. **A.** Serial dilution assay demonstrating the viability of WT and OxyR* overnight cultures after acute exposure to high dose of H_2_O_2_. **B.** Growth curves of WT and OxyR* overnight cultures in standard complex medium (PYE) containing H_2_O_2_ (average ± SD, n = 3) **C.** Catalase activity in WT, OxyR* and Δ*katG* (negative control) strains measured by a cobalt-based activity assay ^18^. The activity of catalase was calculated by normalizing absorbance values to WT (100% activity) and standard (0% activity). Error bars represent the standard deviation (n = 3). *, *P* < 0.05, ***, *P* < 0.001, one-way ANOVA followed by Dunnett’s multiple comparison test to WT. **D.** Serial dilution assay demonstrating the viability of OxyR*, WT, Δ*katG* and OxyR* Δ*katG* overnight cultures after acute exposure to low dose H_2_O_2_. **E.** Left: Western blot showing KatG protein levels in log-and stationary-phase cultures of WT and OxyR* strains (average ± SD, n = 3). Right: quantification of KatG FLAG levels normalized to the loading control (ClpP). The mean of three biological replicates is displayed above its corresponding bar.

### KatG mediates enhanced H_2_O_2_ resistance in the OxyR* strain

To elucidate the mechanism underlying peroxide resistance, we measured H_2_O_2_ degradation using a colorimetric catalase activity assay^18^ (Figure 1C). The oxyR* strain decomposed H_2_O_2_ more rapidly than wild type, indicating elevated catalase activity. Since KatG is the only catalase in *Caulobacter,* we hypothesized it drives the observed H_2_O_2_ resistance. Deletion of *katG* in the OxyR* background substantially reduced cell viability following low-dose H_2_O_2_ treatment, confirming that KatG is responsible for most of the enhanced resistance (Figure 1D). However, OxyR* Δ*katG* cells retained slightly elevated H_2_O_2_ tolerance compared to Δ*katG* alone, suggesting minor contributions from additional factors.

In *Caulobacter*, *katG* expression increases in response to H_2_O_2_ and during entry to stationary phase in an OxyR-dependent manner^14,19,20^. To directly test whether OxyR* constitutively activates *katG*, we measured KatG protein levels by Western blot of a tagged active allele ^21^in logarithmic and stationary phase cultures^_21^. Consistent with expectations, wild type cells showed phase-dependent *katG* induction, with high levels only during stationary phase (Figure 1E). In contrast, the OxyR* strain maintained similarly elevated KatG levels in both phases, indicating constitutive activation nearly tenfold higher than stationary-phase wild type cells. This elevated *katG* expression accounts for the heightened H_2_O_2_ resistance of the oxyR* strain.

### The oxyR* mutation broadly alters the OxyR regulon

To assess whether the A105V substitution affects OxyR targets genome-wide, we performed RNA-seq on log-phase oxyR* cultures. Analysis identified 120 differentially expressed genes with >2-fold change (16 upregulated, 104 downregulated) spanning diverse functional categories (Figure 2A) (Table S1). The most striking upregulation involved the canonical OxyR targets *katG*, *ahpC*, and *ahpF*^14^. The magnitude of *katG* upregulation matched our enzymatic and Western blot data. Notably, *oxyR* itself was upregulated, suggesting the A105V substitution impairs OxyR’s negative autoregulation^14^. Several additional genes of unrelated function were also upregulated, along with a small noncoding RNA downstream of *katG*.

**Figure 2.**
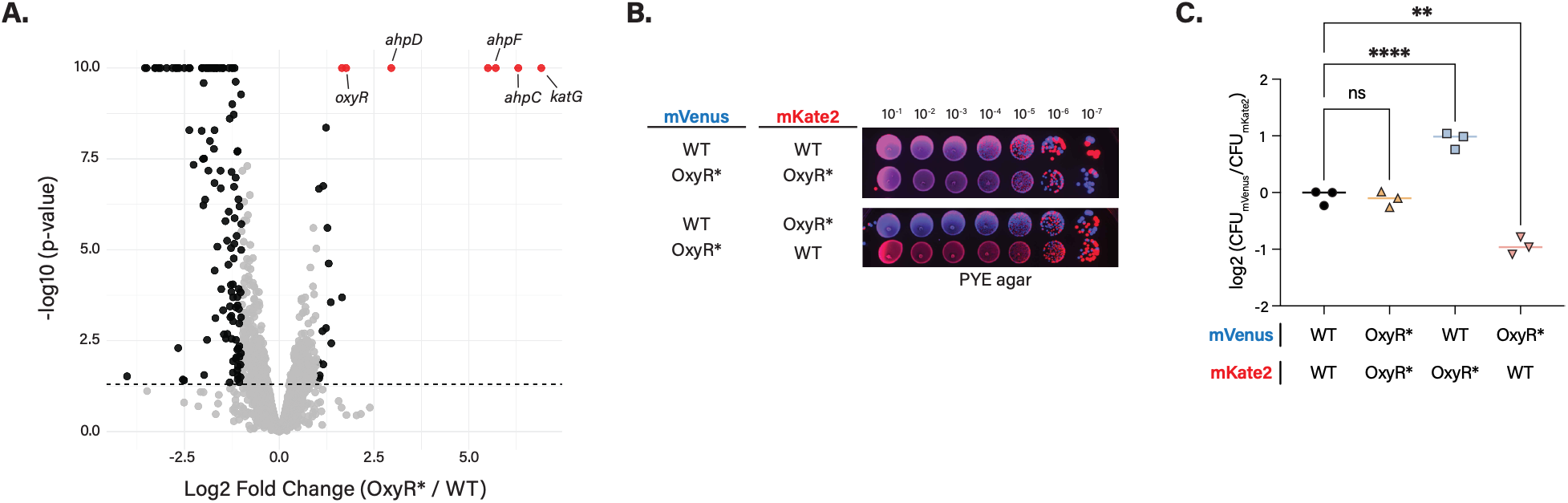
OxyR* strain has growth defect under nonstress conditions. **A.** Volcano plot showing differential gene expression between log-phase cultures of OxyR* and WT (with three biological replicates); gray dots represent all genes, black dots represent differentially regulated genes, and red dots highlight significantly upregulated genes, with –log_10_(p) values capped at 10. Dashed horizontal and vertical lines indicate the significance threshold (p = 0.05) and log2 fold change cutoff (±1), respectively. Select OxyR targets are labeled. **B.** Serial dilution assay of fluorescent protein (mVenus or mKate2)–expressing WT and OxyR* strains cocultured overnight in PYE medium containing kanamycin. **C.** Ratio of mVenus-to mKate2-expressing CFUs of each coculture spread on PYE agar plates. Error bars represent the standard deviation (n = 3). **, *P* < 0.01, ****, *P* < 0.001; one-way ANOVA followed by Dunnett’s multiple comparison test against the mVenus/mKate2 ratio of WT mVenus + WT mKate2 control mixture.

The functional consequences of these expression changes aligned with phenotypic observations. *ahpC* and *ahpF* upregulation correlated with enhanced TBHP resistance (Figure S1), consistent with alkyl hydroperoxide reductase substrate specificity^22^. In contrast, superoxide dismutase expression remained unchanged, explaining the lack of paraquat resistance in OxyR* (Figure S1).

### Constitutive OxyR activation imposes a fitness cost during unstressed growth

Given the transcriptomic changes in OxyR*, we questioned whether activation of the OxyR regulon imposes a fitness cost or advantage during standard growth. Growth curves of monocultures revealed no detectable difference between wild type and OxyR* in either complex (PYE) or defined (M2 glucose) media (Figure 1B, Figure S2A). To detect more subtle effects on fitness, we performed competition assays using strains stably labeled with mVenus or mKate2 (Figure S2B). Control experiments confirmed that reporter expression did not affect oxidative stress resistance (Figure S2C). In co-culture, OxyR* strain exhibited a clear competitive disadvantage after overnight growth (≥12 doublings), regardless of reporter expressed. On serial dilution plates, wild type fluorescence predominated (Figure 2B), and when cocultures were plated on agar, wild type yielded approximately twofold more CFUs than OxyR* (Figure 2C).

To identify the molecular basis of this fitness defect, we tested whether overproduction of the two most highly upregulated enzymes, KatG and AhpCF, accounts for the reduced competitive fitness. We generated OxyR* Δ*katG* and OxyR* Δ*ahpCF* strains and competed each against the parent OxyR* strain under standard growth conditions overnight. Neither deletion rescued the fitness disadvantage (Figure S2D), indicating that KatG and AhpCF overproduction do not drive the competition defect. We concluded that these individual antioxidants were not likely to be the sole determinants of the reduced competitive fitness.

### AhpCF mediates selective sensitivity to reductively activated drugs

Previous work on genes controlled by OxyR during oxidative stress has suggested a potential link to the DNA damage response^14^. We therefore hypothesized that the OxyR* strain may exhibit an altered response to damage induced by genotoxic agents. Viability assays revealed that OxyR* is markedly sensitized to mitomycin C (MMC) (Figure 3A). Notably, sensitivity to other genotoxins (cisplatin, methyl methanesulfonate, 4-nitroquinoline 1-oxide) remained unchanged, indicating selectivity for MMC rather than broad genotoxin hypersensitivity.

**Figure 3.**
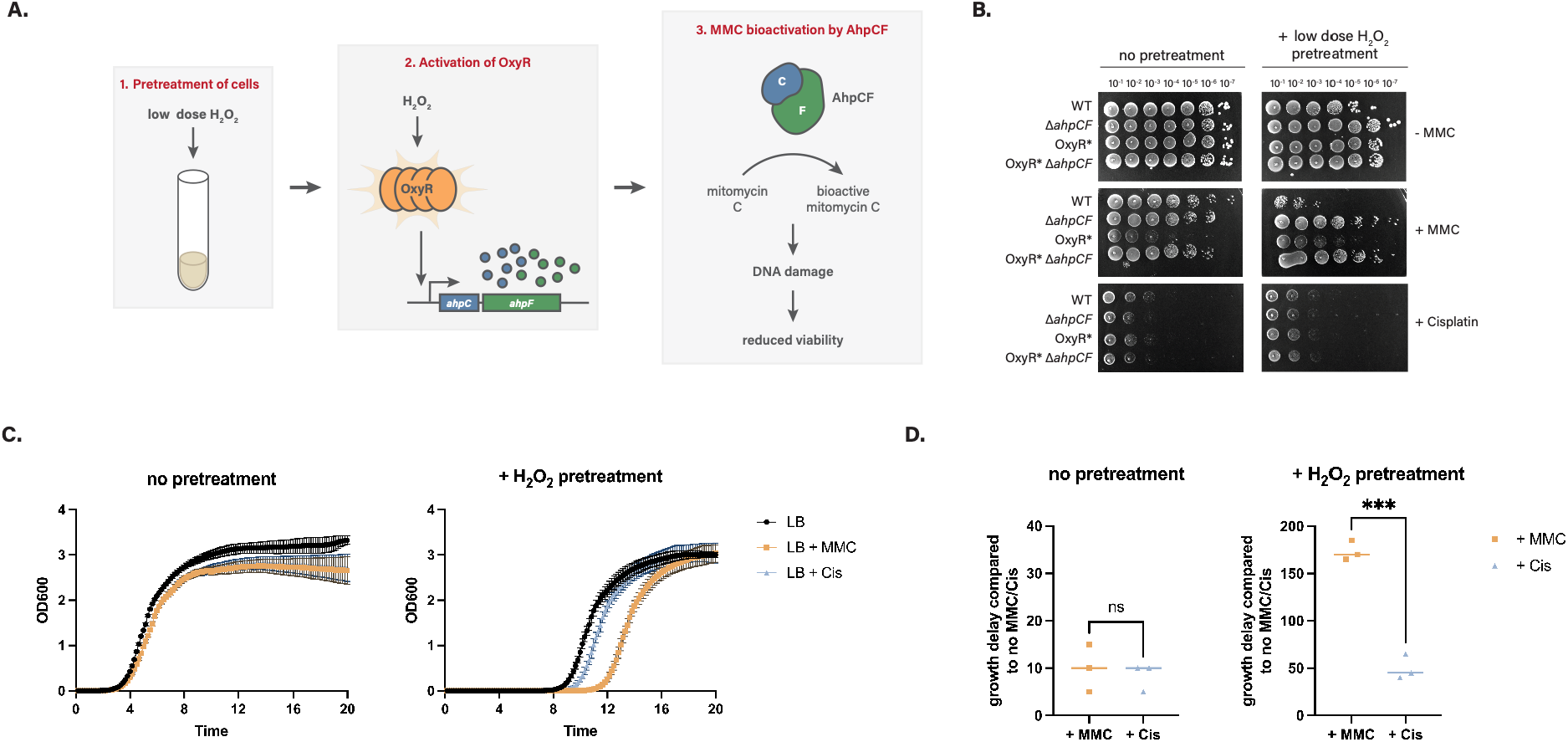
*ahpCF* overexpression in OxyR* strain causes sensitivity to MMC but not to other genotoxic agents. **A.** Serial dilution assays showing viability of WT and OxyR* strains in PYE agar plates containing µg/mL mitomycin C (MMC), methyl methanesulfonate (MMS), cisplatin and 4-nitroquinoline 1-oxide (4NQO). **B.** Serial dilution assay of WT, OxyR*, Δ*ahpCF*, and OxyR* Δ*ahpCF* strains on PYE agar plates ± 0.5 µg/mL MMC. **C. Top**, schematic representation of the xylose-inducible *ahpCF* overexpression vector used. **Bottom**, viability of Δ*ahpCF* and OxyR* Δ*ahpCF* strains transformed with either the empty vector (EV) or xylose-inducible *ahpCF* overexpression vector (*ahpCF*++) on PYE agar plates containing 0.2% xylose ± 0.125 µg/mL MMC. **D.** Growth curves of WT, OxyR* and OxyR* Δ*ahpCF* overnight cultures in PYE medium ± nitrofurantoin (NIT). **E.** Graphs showing the time it takes for OxyR* and OxyR* Δ*ahpCF* cells to reach an OD600 of 1 in PYE media ± NIT compared to WT control (n = 3 biological replicates). **, P < 0.005; Welch’s t-test. Horizontal lines are median values.

Mitomycin C requires reductive activation and can be processed by multiple cellular reductases^23–31^. Given that the OxyR* strain overproduces several reductases, we reasoned that altered reductase expression might account for MMC hypersensitivity. To identify the responsible factor, we deleted the *ahpCF* operon, which encodes the most upregulated reductase. Strikingly, loss of *ahpCF* fully restored MMC resistance in the OxyR* strain (Figure 3B), implicating AhpCF as the determinant of heightened MMC sensitivity. Next, to confirm whether AhpCF overexpression is sufficient for MMC sensitivity, we constructed a plasmid enabling xylose-inducible *ahpCF* expression. MMC sensitivity substantially increased upon elevation of AhpCF (Figure 3C), which we attributed to increased reduction of MMC to its activated form. Consistent with this interpretation, purified AhpC is able to reduce MMC *in vitro* (Figure S3A). Together, these results suggests that induction of the AhpCF reductase during oxidative stress results in a collateral consequence of MMC activation.

To assess whether the AhpCF-dependent vulnerability extends beyond MMC, we tested additional reductively activated compounds. The OxyR* strain showed modest but reproducible increased sensitivity to nitrofurantoin (NIT), a prodrug whose reduction is primarily catalyzed by NfsA and NfsB^32,33^ in *Escherichia coli*, with reported involvement of AhpF activity^34^. Deletion of *ahpCF* in the OxyR* background partially restored NIT resistance (Figure 3D, E), suggesting that AhpCF contributes to NIT activation, although additional reductases likely participate. By contrast, OxyR* showed no increased sensitivity to 4-nitroquinoline 1-oxide (Figure 3A), indicating that enzymes responsible for 4NQO reduction are not upregulated in the mutant. Collectively, these findings support a model in which AhpCF possesses sufficient broad reductase activity to increase the pool of active MMC and NIT upon their entry into the cell.

To establish functional specialization within the OxyR regulon, we verified that *ahpCF* deletion does not compromise H_2_O_2_ resistance and found that OxyR* Δ*ahpCF* retained full resistance to acute, high-dose H_2_O_2_ exposure (Figure S3B). Conversely, *katG* deletion had no effect on MMC sensitivity in the OxyR* background (Figure S3C). These results demonstrate a clear division of labor where *katG* upregulation specifically mediates enhanced H_2_O_2_ resistance, while *ahpCF* upregulation specifically sensitizes cells to reductively activated prodrugs.

### Induction of *ahpCF* by external oxidants results in collateral sensitivity to MMC

Since *ahpCF* overexpression in the OxyR* strain confers MMC sensitivity, we predicted that H_2_O_2_-mediated OxyR activation would similarly induce *ahpCF* expression in wild type cells and sensitize them to MMC (Figure 4A). Log-phase cultures transiently exposed to low-dose H_2_O_2_ were subsequently assessed for cell viability on plates containing MMC or cisplatin (Figure 4B). H_2_O_2_ pretreatment significantly increased MMC sensitivity in wild type cells but had no effect on Δ*ahpCF* cells. Notably, H_2_O_2_ pretreatment did not increase cisplatin sensitivity in either strain, demonstrating that *ahpCF*-dependent activation is selective for MMC. As expected, the OxyR* strain exhibited maximal MMC sensitivity without further sensitization by H_2_O_2_, consistent with its constitutively elevated *ahpCF* expression. Together, these findings reveal that activation of oxidative stress defenses creates collateral sensitivity to reductively activated drugs through AhpCF-dependent bioactivation.

**Figure 4.**
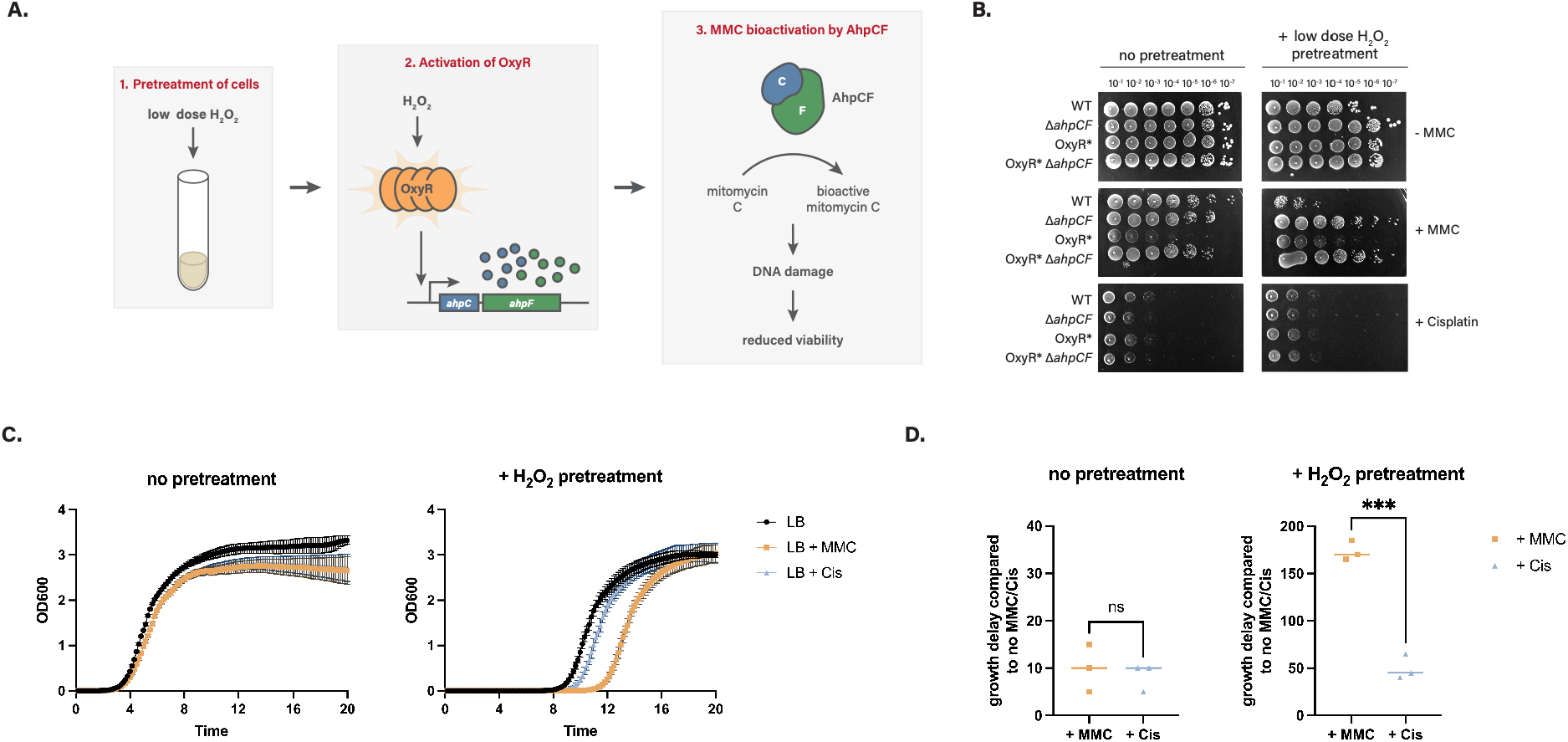
Induction of *ahpCF* by external oxidants results in collateral sensitivity to MMC. **A.** Schematic cartoon illustrating the experimental plan and the working hypothesis. **B.** Serial dilution assays showing viability of H_2_O_2_ treated or untreated mid-log cultures of WT, Δ*ahpCF*, OxyR* and OxyR* Δ*ahpCF* strains on PYE agar plates containing MMC or Cisplatin. **C.** Growth curves of H2O_2_ treated or untreated *E*. *coli* K12 mid-log cultures in LB media supplemented with 0.75 µg/mL MMC or 25 µg/mL cisplatin. **D.** Graphs showing the time delay (in minutes) for H_2_O_2_ treated or untreated *E*. *coli* K12 mid-log cultures to reach an OD600 of 1 in LB media containing MMC or Cisplatin compared to no MMC/Cis control (n = 3 biological replicates). ***, P < 0.005; Welch’s t-test. Horizontal lines are median values.

To determine if this phenomenon is conserved across bacterial species, we exposed log-phase *Escherichia coli* K-12 to brief H_2_O_2_ treatment and assessed drug sensitivity (Figure 4C). Without pretreatment, MMC and cisplatin caused similar growth delays. However, H_2_O_2_ pretreatment had a significant effect on MMC sensitivity (16x delay in growth) with less effect on cisplatin (5x delay in growth), mirroring our results in *Caulobacter*. These findings in divergent bacterial species demonstrate that collateral sensitivity to reductively activated drugs might represent a conserved consequence of antioxidant responses.

## DISCUSSION

Antioxidant defense systems face a fundamental tension because they must provide rapid, effective protection from ROS while minimizing costs when stress is absent. Our findings demonstrate that activation of the OxyR-controlled program in *Caulobacter crescentus* exemplifies this trade-off. A single gain-of-function *oxyR* allele confers strong peroxide resistance through elevated catalase-peroxidase (KatG) expression yet simultaneously creates selective vulnerability to the reductively activated genotoxin mitomycin C (MMC) via AhpCF overproduction and reduces competitive fitness under nonstress conditions (Figure 5).

**Figure 5.**
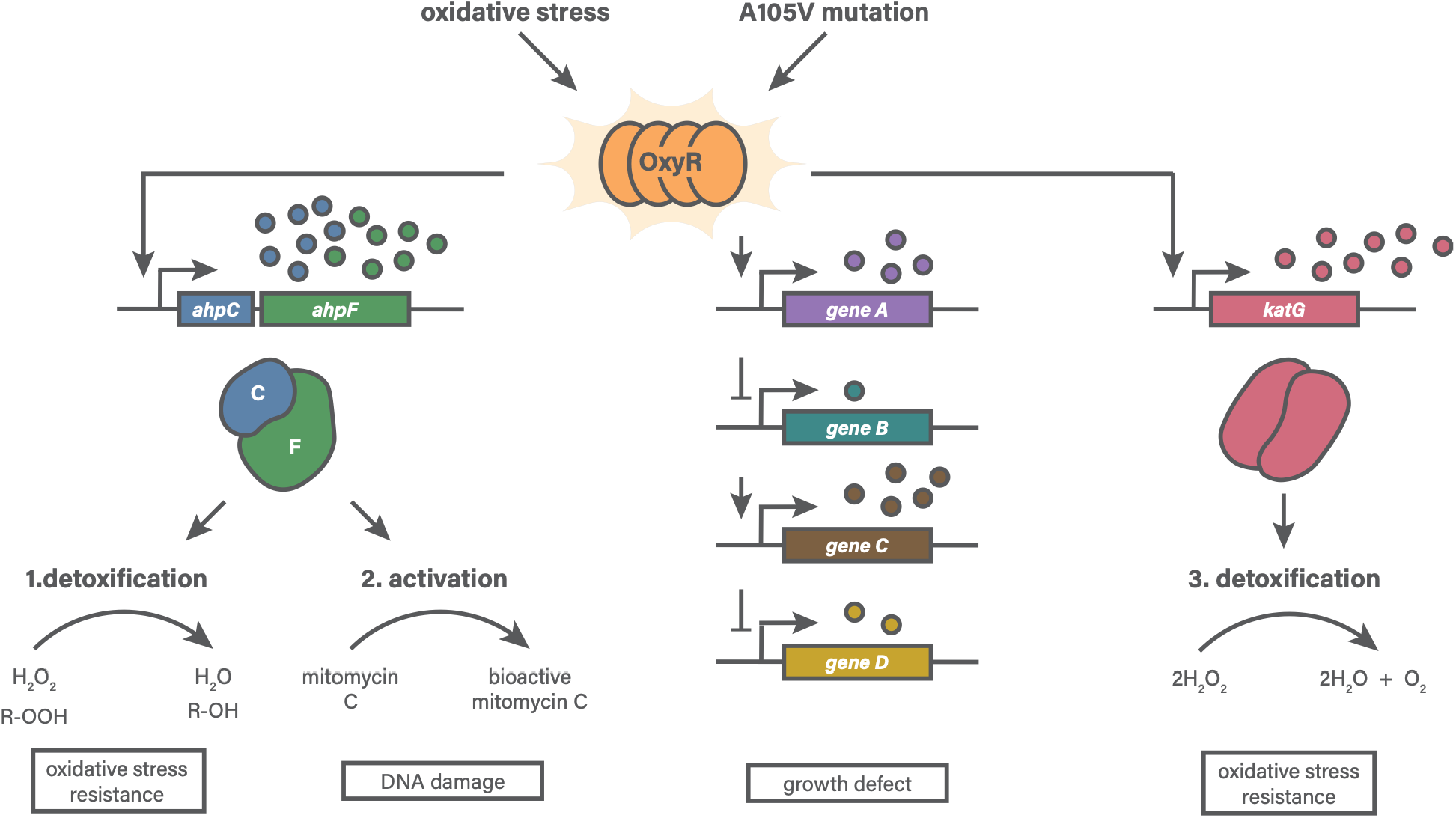
Model for OxyR-dependent regulation of antioxidant genes and its physiological consequences. Schematic illustrating how OxyR is activated either by oxidative stress (elevated H_2_O_2_) or by the A105V mutation, which likely locks OxyR in an active conformation. Activated OxyR induces transcription of antioxidant genes, including katG, ahpC, ahpF, and additional members of the OxyR regulon. Increased KatG and AhpCF levels enhance detoxification of H_2_O_2_, resulting in elevated peroxide resistance. However, high AhpCF expression also creates a collateral vulnerability, sensitizing cells to mitomycin C (MMC).

The competitive disadvantage of the OxyR* strain illuminates why redox-sensitive regulators must remain under stringent control. We initially hypothesized that overproduction of the two most highly upregulated enzymes, AhpCF or KatG, might account for the fitness defect. However, deletion of either gene individually failed to rescue competitive fitness in the OxyR* background, indicating that enzyme overproduction alone does not drive the disadvantage. This result suggests that the broader metabolic burden of sustained upregulation across over many differentially regulated genes in the OxyR* mutant collectively imposes costs that reduce growth efficiency. Notably, beyond this general regulatory burden, our work reveals a more specific and mechanistic fitness cost as constitutive OxyR activation sensitizes cells to reductively activated genotoxins like MMC through AhpCF-dependent drug bioactivation.

This AhpCF-dependent bioactivation reflects a fundamental property of reductases where their inherent promiscuity in substrate recognition enables them to bioactivate structurally diverse prodrugs^29,35,36^. This principle is not unique to bacteria. In eukaryotic cells, aberrant activation of the antioxidant regulator Nrf2 through KEAP1 suppression is leads to increased NQO1 and CYPOR reductase expression and thought to lead to enhanced MMC reduction^36^. Our work demonstrates that *Caulobacter* employs an analogous mechanism: when upregulated during antioxidant responses, elevated AhpCF reductase activity simultaneously enhances activation of certain antimicrobials, creating selective vulnerability to reductively activated drugs. This specificity reflects the substrate preferences of AhpCF and demonstrates that the trade-off is mechanistically defined rather than a general consequence of stress activation.

Transient H_2_O_2_ exposure recapitulates this MMC sensitization in both *Caulobacter* and in *Escherichia coli*, suggesting that it might emerge as a general consequence of peroxide-inducible reductase upregulation across divergent bacterial species. However, whether this trade-off is broadly conserved across all bacterial lineages remains an open question. The enzymatic determinants of prodrug activation differ substantially among bacteria, reflecting differences in enzyme expression patterns, substrate affinities, and regulatory architecture. Consequently, both the magnitude and specificity of collateral sensitivity will likely depend on the particular enzyme repertoire and regulatory context of each organism.

## MATERIALS AND METHODS

### Growth media and conditions

All *Caulobacter crescentus* strains were grown in peptone-yeast extract (PYE) medium or in M2 glucose defined medium. PYE medium: 2 g/L peptone, 1 g/L yeast extract, 1 mM MgSO_4_, and 0.5 mM CaCl_2_. M2-Glucose medium: 0.5 mM MgSO_4_, 0.5 mM CaCl_2_, 0.01 mM ferrous sulfate chelate solution (Sigma Aldrich), 0.2% (w/v) glucose and autoclaved 1x M2 salts. 20x M2 salts solution: 17.4 g/L Na_2_HPO_4_, 10.6 g/L KH_2_PO_4_, and 5 g/L NH_4_Cl. For growth on solid media, PYE media containing 1.5% agar (w/v) was used. *Escherichia coli* Top10 and W3110 (K12) strains were cultivated in LB media (10 g/L tryptone, 10 g/L NaCl, 5 g/L yeast extract) and for solid growth, LB media containing 1.5% agar (w/v) was prepared. Gene expression from xylose promoter was induced by adding 0.2% (w/v) xylose prior to viability assays.

The following antibiotic concentrations were used: for solid medium, kanamycin at 25 µg/mL for *Caulobacter crescentus* and 50 µg/mL for *E*. *coli*; gentamycin at 5 µg/mL for *Caulobacter* and 40 µg/mL for *E*. *coli*. For liquid medium, kanamycin was used at 5 µg/mL for *Caulobacter* and 50 µg/mL for *E*. *coli;* gentamycin was used at 0.25 µg/mL for *Caulobacter* and 40 µg/mL for *E*. *coli*; ampicillin was used at 50 µg/mL for *E*. *coli*.

### Plasmids construction

Please refer to Table S2 for detailed information on plasmids and strains. To generate a *Caulobacter crescentus* strain expressing the OxyR* mutant, we amplified a ∼3 kb DNA fragment encompassing the *oxyR* coding sequence along with 1 kb upstream (5′ UTR) and 1 kb downstream (3′ UTR) regions from the original mutant strain. The amplified fragment was inserted into the HindIII–EcoRI sites of the suicide vector pNPTS138 using Gibson assembly, replacing the corresponding region of the vector. The assembly mixture was transformed into *E*. *coli* Top10 (Invitrogen), and colonies carrying the complete construct (pNPTS138-*OxyR\**) were selected on LB agar plates containing kanamycin. For construction of the *ahpCF* deletion plasmid, the 5′ UTR of *ahpC* and the 3′ UTR of *ahpF* were PCR-amplified and assembled with a gentamycin resistance cassette replacing the *ahpCF* coding region. These fragments were cloned into the pNPTS138 vector by Gibson assembly, and the mixture was transformed into *E*. *coli* Top10 cells. Colonies carrying the full plasmid (pNPTS138-Δ*ahpCF*::*gent*) were selected on LB agar containing kanamycin. For xylose-inducible overexpression of *ahpCF*, the coding sequences of *ahpC* and *ahpF* were amplified using primers with 5′ and 3′ overhangs containing NdeI and ApaI restriction sites, respectively. The pBX-MCS-2 plasmid was digested with NdeI and ApaI, and the inserts were ligated into the vector by restriction cloning. Ligation mixtures were transformed into *E*. *coli* Top10 cells, and recombinant colonies were selected on LB agar containing kanamycin. For protein expression, the *Caulobacter ahpC* gene (CCNA_03012) was synthesized by Twist Biosciences as a DNA insert and cloned into the pET23b expression vector via Gibson assembly. The Gibson assembly mixture containing the plasmid (pET23b-ahpC-6xHis) was chemically transformed into competent *E*. *coli* Top10 cells for plasmid amplification and into *E*. *coli* T7 Express cells for protein production.

### Strain construction

All the *Caulobacter crescentus* strains utilized in this study originated from the NA1000 strain. NA1000 *katG*::*M2-FLAG* and *katG*::*M2-FLAG* OxyR* strains were constructed by two-step homologous recombination using plasmids pNPTS138-*katG*::*M2-FLAG* (a gift from Sean Crosson) and pNPTS138-*OxyR\**, respectively. The *katG*::*M2*-*FLAG* Δ*ahpCF*::*gent* strain was generated using the pNPTS138-Δ*ahpCF*::*gent* plasmid. Wild-type or *katG*::*M2*-*FLAG* cells were transformed with the respective pNPTS138 constructs via electroporation and plated on PYE agar containing kanamycin (25 µg/mL) for primary selection. Single colonies were restreaked onto PYE agar supplemented with 3% (w/v) sucrose to select for secondary recombination. Sucrose-resistant colonies were replica-patched onto PYE plates with and without kanamycin; kanamycin-sensitive isolates were considered to have successfully excised the plasmid backbone. For *ahpCF* deletion, colonies were additionally patched onto PYE agar containing gentamycin to confirm insertion of the *gent* cassette. Insertion of the M2-FLAG tag at the *katG* C-terminus using was verified by Western blotting using an anti-FLAG antibody. Replacement of the wild type *oxyR* allele with the A105V mutant, as well as *ahpCF* deletion, were confirmed by whole-genome sequencing. OxyR* Δ*ahpCF* and OxyR* Δ*katG* strains were constructed by transducing the *ahpCF*::*gent* or *katG*::*gent* alleles into the OxyR* background using the φCR30 phage. A previously generated Δ*katG*::*gent* strain was used as a control and as the donor for phage transduction into the OxyR* background^37^. For inducible overexpression of *ahpCF,* the pBX-MCS-2-*ahpCF* plasmid was introduced into NA1000 cells by electroporation, and transformants were selected on PYE agar containing kanamycin. Fluorescent protein (mVenus or mKate2) expressing strains were generated via phage transduction and colonies were selected on PYE agar plates containing kanamycin.

### Whole genome sequencing

Genomic DNA from overnight cultures were isolated using Monarch^®^ Genomic DNA Purification Kit (New England Biolabs) and Nanodrop One/One^C^ spectrophotometer (Thermo Fisher) was used to determine concentration of DNA. Isolated genomic DNA samples were sent to SeqCenter, LLC (MiGS - Microbial Genome Sequencing Center, Pittsburgh, PA) for Illumina whole genome sequencing, or to Plasmidsaurus for bacterial genome sequencing using Oxford Nanopore Technology with custom analysis and annotation. Breseq was used to analyze raw FASTQ files to confirm gene deletions and substitutions^38^.

### Bacterial growth curves

For growth curves, 100 µM hydrogen peroxide, 75 µM tert-butyl hydroperoxide (TBHP), 50 µM paraquat (PQ) or 4 µg/mL nitrofurantoin (NIT) were added to growth media. Overnight cultures were diluted to an OD600 of 0.1 in PYE medium. A 4 µL aliquot of each diluted culture was added to 196 µL of growth medium in sterile, clear, flat-bottom 96-well plates (Fisher Scientific). Plates were incubated at 30 °C in BioTek Epoch 2 microplate reader with continuous shaking at 567 cpm, and absorbance measurements at 600 nm were recorded every 20 minutes for 24 hours.

### H_2_O_2_ cell viability experiments

Acute H_2_O_2_ treatment conditions are as follows: Overnight cultures treated with 10 mM (low dose) or 100 mM (high dose) for 15 minutes. Treated cultures were centrifuged for 2 minutes at 6000xg and the supernatant was removed. Fresh media was added to tubes, and cell pellets were gently resuspended. OD600 of cultures were measured and normalized to OD600 of 0.1. Treated and untreated overnight cultures were then serially diluted and 3 µL of each diluent were spotted to PYE agar plates. Treated or untreated mid-log cultures were spotted on PYE agar plates containing MMC or cisplatin. Spotted plates were incubated at 30 °C for 2 days.

### Determination of catalase activity

H_2_O_2_ levels were determined using colorimetric assay described in Hadwan et al, 2018^18^. Overnight cultures were normalized to OD600 = 1 in 5 mL PYE and centrifuged at 6000xg for 5 minutes at room temperature and resuspended in 500 µL phosphate buffer (pH 7.0, 50 mM). 10 mM H_2_O_2_ was added to cell suspensions, and the standard (except blank) and tubes were incubated at 30 °C for 2 minutes. 6 mL working solution was added to each tube and tubes were briefly vortexed prior to incubation at room temperature in the dark for 15 minutes. The tubes containing cultures were centrifuged at 21000xg for 3 minutes and the absorbance of supernatant at 440 nm was measured. The absorbance values were compared with blank and standard to assess catalase activity.

### Western Blotting

WT and OxyR* stationary phase cultures were normalized to OD600 = 1 in 1 mL PYE and mid-log phase cultures (OD600 ∼ 0.5) were normalized to OD600 = 0.5 in 2 mL of PYE. Normalized samples were centrifuged at >15000xg for 2 minutes at room temperature. Pellets were resuspended in Laemmli buffer (100 mM Tris pH 6.8, 4% (w/v) SDS, 20% (v/v) glycerol, 0.2% (w/v) bromophenol blue, 10% (v/v) ß-mercaptoethanol). Samples were incubated at 95 °C for 15 minutes and centrifuged at >15000xg for 10 minutes. Supernatants were electrophoresed in 10% Bis-Tris polyacrylamide gels casted using SureCast Gel Handcast System (Thermo Fisher). The gels were transferred to nitrocellulose membranes (Cytiva) at 10V for 1 hour using mini blot module (Thermo Fisher). The membranes were blocked in 5% milk (w/v) in TBS-T and incubated overnight with anti-FLAG (Sigma-Aldrich) or anti-ClpP primary antibodies. For secondary antibody incubation, fluorescent goat anti-rabbit (680LT) or goat anti-mouse (800CW) antibodies (LiCor) were used. The membranes were imaged on LiCor instrument using ImageStudio software.

### RNA-Seq sample preparation

Overnight cultures of WT and OxyR* strains were backdiluted to OD600 = 0.05 and grown to mid-log phase (OD600 ∼ 0.5). The cultures were normalized to an OD600 of 0.5 in 2 mL of PYE media and centrifuged at 21000xg for 2 minutes at room temperature. Pellets were then frozen in liquid nitrogen and sent to SeqCenter, LLC (MiGS - Microbial Genome Sequencing Center, Pittsburgh, PA) for total RNA extraction followed by RNA sequencing with rRNA depletion using custom probes. Raw FASTQ files were analyzed using a pipeline inspired by BactSeq (Adam Dinan, 2022).

### Competition assays

mVenus-or mKate2-expressing WT and OxyR* strains were grown overnight in PYE medium supplemented with kanamycin. Kanamycin was included in all media for competition experiments with fluorescent strains. Overnight cultures were backdiluted and grown to mid-log phase, then normalized to OD600 = 0.001 in fresh medium and co-cultured with their respective competitor strains overnight at 30 °C. Each pair was cocultured for ≥ 12 doublings. OD600 values of each coculture were measured and normalized to OD600 = 0.1 in 1.5 mL tubes. Normalized cultures were transferred to sterile 96-well plates for serial dilution. 3 µL from each diluent were spotted on PYE agar plates containing kanamycin. For CFU determination of competing strains, 50 µL of the 10^-6^ diluent of the coculture mixtures were spread on PYE agar plates supplemented with kanamycin. All plates were incubated at 30 °C for 2 days and resulting CFUs were counted.

### DNA damage experiments

Overnight cultures of *Caulobacter* strains grown in PYE media were normalized according to OD600 values. Cultures were serially diluted and 3 µL from each dilution was spotted onto PYE agar plates containing DNA damaging agents at following concentrations: mitomycin C (MMC) at 0.25 or 0.5 µg/mL, methyl methanesulfonate (MMS) at 1.18 mM, cisplatin at 2.5 µg/mL and 4-Nitroquinoline 1-oxide (4NQO) at 0.8 µg/mL. Spotted plates were incubated at 30 °C for 2 days.

### *ahpCF* overexpression viability assays

Strains carrying either the empty vector (EV; pBX-MCS-2) or the xylose-inducible *ahpCF* expression plasmid (*ahpCF*++; pBX-MCS-2-*ahpCF*) were grown overnight in PYE medium containing kanamycin. Six hours before plating, xylose was added to each culture to a final concentration of 0.2% (w/v). After induction, OD600 values were measured and normalized to 0.1. Normalized cultures were transferred to sterile 96-well plates for serial dilution, and 3 µL of each dilution (10^-1^ to 10^-7^) were spotted onto PYE agar plates containing 0.2% xylose with or without 0.125 µg/mL MMC.

### H_2_O_2_-dependent MMC viability experiments

Mid-log *Caulobacter* cultures were treated with 10 mM H_2_O_2_ for 1 hour at 30 °C. Mid-log *E*. *coli* W3110 cultures were treated with 20 mM H_2_O_2_ for 30 minutes at 37 °C. Treated cultures were centrifuged for 2 minutes at 6000xg and the supernatant was removed. Pellets were resuspended in fresh media and OD600 measurements were taken. For *Caulobacter*, treated and untreated cells were then serially diluted and 3 µL of each diluent were spotted on PYE agar plates containing 0.25 µg/mL MMC or 2.5 µg/mL cisplatin. Spotted plates were incubated at 30 °C for 2 days.

For *E*. *coli,* treated and untreated cells were diluted and transferred to 96-well plates with wells containing LB ± 0.75 µg/mL MMC or 25 µg/mL cisplatin with a starting OD600 = 0.005. Plates were placed in BioTek Epoch 2 microplate reader with continuous shaking at 567 cpm, and absorbance measurements at 600 nm were recorded every 5 minutes for 36 hours at 37 °C.

### Purification of *Caulobacter* AhpC protein

*E*. *coli* T7 Express cells containing the pET23b-ahpC-6xHis plasmid were grown overnight in 20 mL LB medium supplemented with ampicillin at 37°C. The overnight culture was diluted into 2 L LB containing ampicillin and grown at 37°C until OD600 reached 0.6. Protein expression was induced by addition of IPTG to a final concentration of 0.4 mM. The temperature was reduced to 20°C and cultures were grown overnight. Cells were harvested by centrifugation, and the pellet was resuspended in 20 mL lysis buffer (50 mM Tris pH 8.0, 300 mM NaCl, 10% glycerol, 10 mM imidazole). The resuspended cells were lysed using a microfluidizer (Microfluidics, Newton, MA), and the lysate was clarified by centrifugation at 14,000xg for 30 minutes at 4°C. The supernatant was loaded onto pre-equilibrated gravity-flow Ni-NTA resin. The resin was washed with lysis buffer and eluted with elution buffer (50 mM Tris pH 8.0, 300 mM NaCl, 10% glycerol, 300 mM imidazole). The eluted protein was mixed with Ulp1-His protease and dialyzed overnight into lysis buffer without imidazole to enable 6xHis-SUMO tag cleavage and buffer exchange. The cleavage mixture was loaded onto gravity-flow Ni-NTA resin to separate cleaved AhpC from uncleaved protein and Ulp1 protease.

### In vitro MMC reduction experiments

To assess AhpC-mediated MMC bioactivation in vitro, we performed a spectrophotometric reduction assay. Mitomycin C (MMC) reduction was assessed by monitoring changes in UV-visible absorbance spectra. AhpCF catalyzes substrate reduction through a catalytic cycle in which the AhpF subunit regenerates reduced AhpC from oxidized AhpC using NADH as an electron donor. To mimic this catalytic cycle and enable AhpC to reduce MMC, we substituted the AhpF/NADH electron donation system with dithiothreitol (DTT), a chemical reducing agent capable of directly reducing the active site cysteines of AhpC. Reaction mixtures (250 µL) containing 50 µM MMC in sodium phosphate buffer (50 mM, pH 6.8) were prepared with the following additions: (1) no additions (MMC control), (2) 1 mM DTT, (3) 5 mM DTT, (4) 10 µM purified AhpC, (5) 10 µM bovine serum albumin (BSA), (6) 10 µM AhpC plus 1 mM DTT, (7) 10 µM BSA plus 1 mM DTT, (8) 10 µM AhpC plus 5 mM DTT, and (9) 10 µM BSA plus 5 mM DTT. All mixtures were incubated at 37°C for 1 hour, and absorbance spectra (300-700 nm) were measured using a spectrophotometer. MMC reduction was quantified by monitoring absorbance changes at 365 nm, with decreased absorbance indicating reduction of MMC.

## ACKNOWLEDGEMENTS

This project was supported by NIH R35GM130320 to P.C. K.Y. was supported in part through the University of Massachusetts Amherst NIH Chemistry-Biology Interface Training Program (NIH T32GM139789). A.C was a CAFE program summer scholar at UMass Amherst. We thank Chien lab members for helpful discussion.

## AUTHOR CONTRIBUTIONS

Conceptualization: K.Y. and P.C.; investigation: K.Y. and A.C.; writing: K.Y. and P.C.; funding acquisition: P.C.; supervision: P.C.

---

**Supplementary Figure 1, related to Figure.**
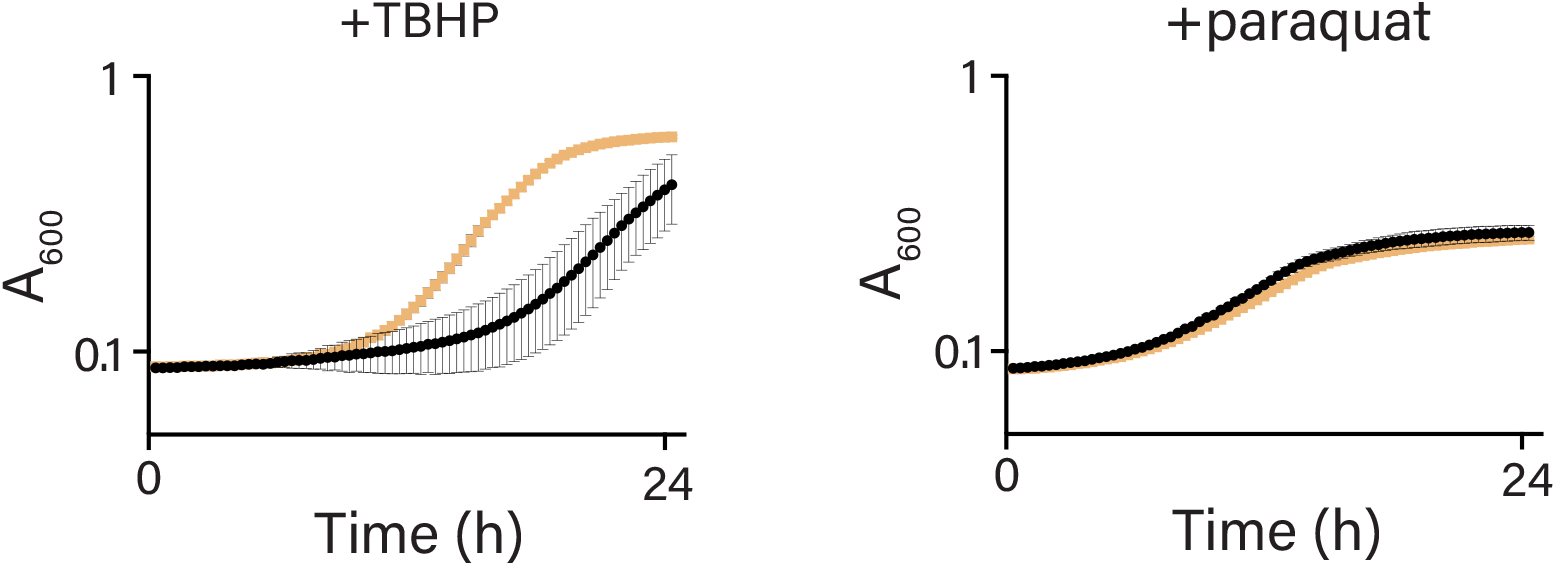
Growth curves of WT (black) and OxyR*(orange) overnight cultures in standard complex medium (PYE) containing tert-butyl hydroperoxide (TBHP) or paraquat (PQ) respectively. Error bars represent standard deviation (n = 3).

**Supplementary Figure 2, related to Figure 2.**
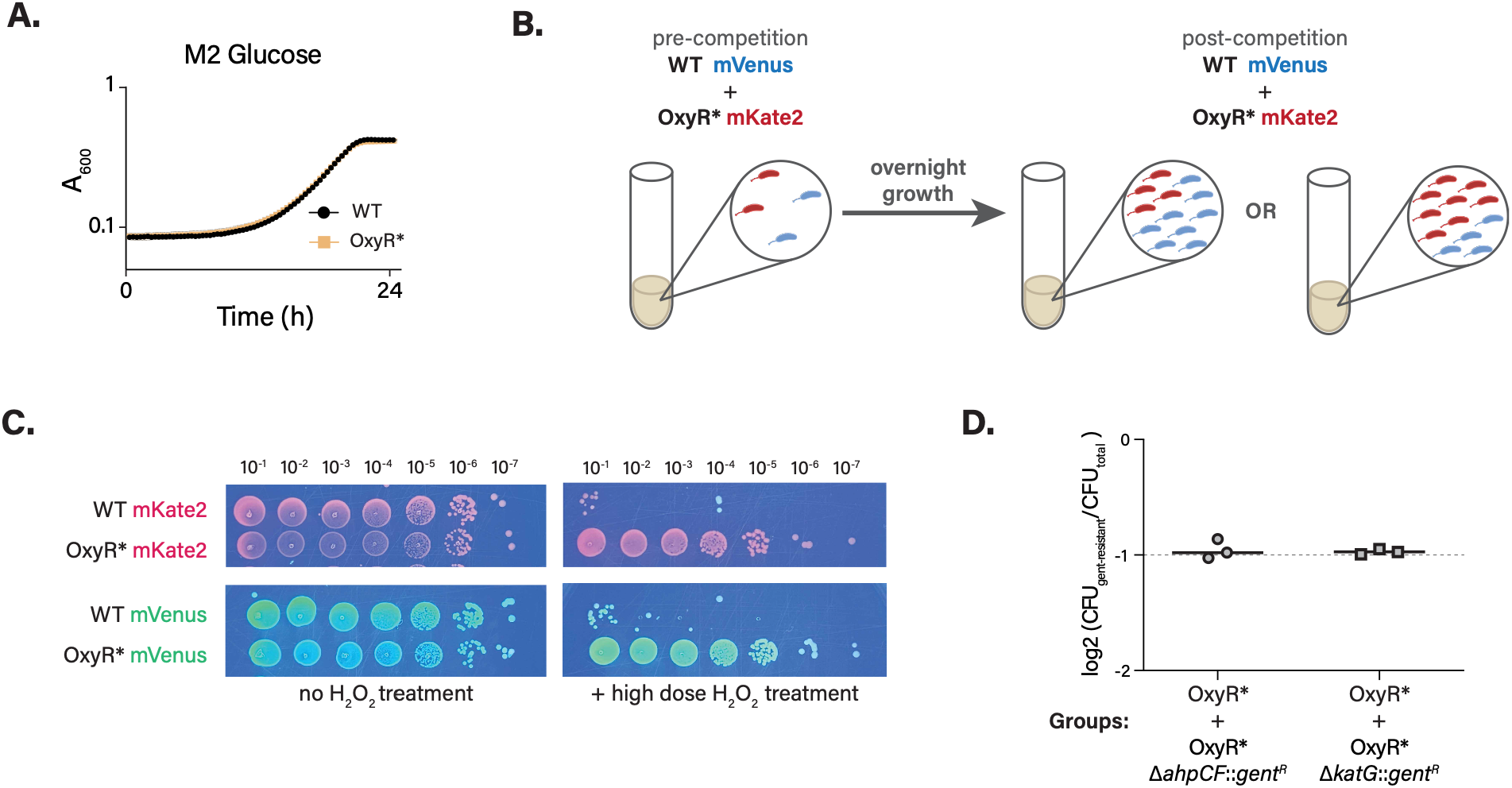
**A.** Growth curves of WT and OxyR* overnight cultures in M2 glucose medium (n = 3). **B.** Schematic cartoon illustrating the competition assay between WT and OxyR* strains. **C.** Serial dilution assay of fluorescent protein (mVenus or mKate2)–expressing WT, and OxyR* strains on PYE agar (left), on MMC-containing PYE agar (center) and on PYE agar plates following treatment with high dose of H_2_O_2_ (right). **D.** Ratio of gentamycin-resistant CFUs to total CFUs of each coculture spread on PYE agar plates. Horizontal lines are median values.

**Supplementary Figure 3, related to Figure 3.**
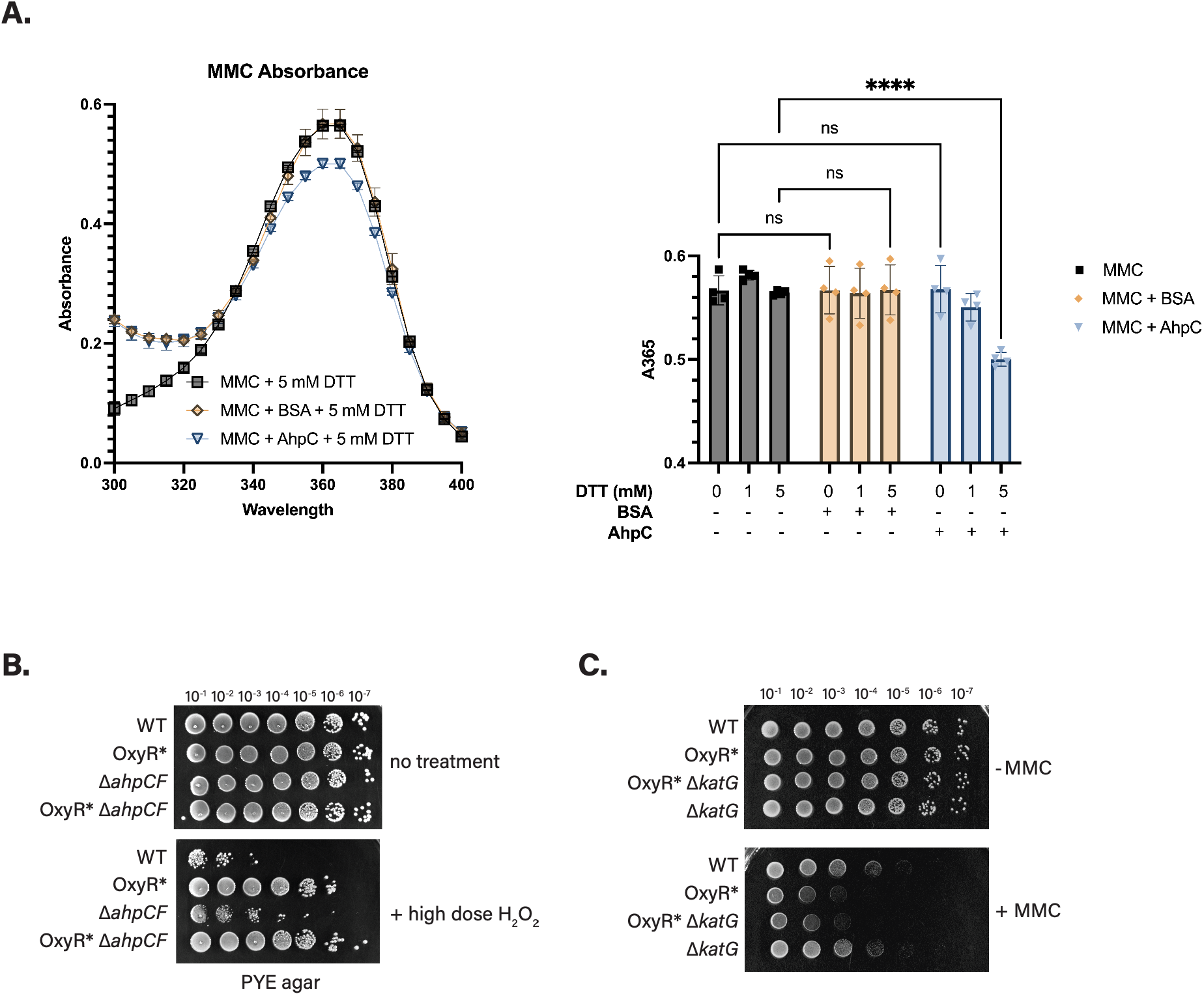
**A.** Left, Absorbance spectra of 50 µM MMC incubated with DTT, purified AhpC (10 µM), or BSA (10 µM), alone or in the indicated combinations. Right, quantification of absorbance at 365 nm (A365) from the spectra shown in the left panel. Error bars represent the standard deviation (n = 4). ***, *P* < 0.0005, two-way ANOVA followed by Sidak’s multiple comparison test to MMC ± DTT controls. **B.** Serial dilution assays of WT, OxyR*, Δ*ahpCF*, and OxyR* Δ*ahpCF* strains on PYE agar plates following treatment with high dose of H_2_O_2_. **C.** Serial dilution assay of WT, OxyR*, OxyR* Δ*katG*, and Δ*katG* strains on PYE agar ± MMC.

